# Terminally differentiated effector memory T cells associate with cognitive and AD-related biomarkers in an aging-based community cohort

**DOI:** 10.1101/2023.11.27.568812

**Authors:** Edric Winford, Jenny Lutshumba, Barbara J. Martin, Donna M. Wilcock, Gregory A. Jicha, Barbara S. Nikolajczyk, Ann M Stowe, Adam D. Bachstetter

**Author notes:** Corresponding author Correspondence to: Adam D. Bachstetter Full address: 741 S. Limestone St. Rm B459 Lexington, KY 40536, USA. **Abbreviations:** Alzheimer’s Disease (AD); Alzheimer’s disease and related dementia (ADRD); peripheral blood mononuclear cells (PBMCs); terminally differentiated memory T cells (TEMRAs); central memory T cells (TCM); effector memory T cells (TEM); β amyloid (Aβ); neurofilament light (Nf-L); glial fibrillary acidic protein (GFAP); clinical dementia rating scale (CDR); Mini-Mental State Examination (MMSE); Montreal Cognitive Assessment (MoCA).

## Abstract

**Background and Purpose:** The immune response changes during aging and the progression of Alzheimer’s disease (AD) and related dementia (ADRD). Terminally differentiated effector memory T cells (called T_EMRA_) are important during aging and AD due to their cytotoxic phenotype and association with cognitive decline. However, it is not clear if the changes seen in T_EMRAs_ are specific to AD-related cognitive decline specifically or are more generally correlated with cognitive decline. This study aimed to examine whether T_EMRAs_ are associated with cognition and plasma biomarkers of AD, neurodegeneration, and neuroinflammation in a community-based cohort of older adults.

**Methods:** Study participants from a University of Kentucky Alzheimer’s Disease Research Center (UK-ADRC) community-based cohort of aging and dementia were used to test our hypothesis. There were 84 participants, 44 women and 40 men. Participants underwent physical examination, neurological examination, medical history, cognitive testing, and blood collection to determine plasma biomarker levels (Aβ42/Aβ40 ratio, total tau, Neurofilament Light chain (Nf-L), Glial Fibrillary Acidic Protein (GFAP)) and to isolate peripheral blood mononuclear cells (PBMCs). Flow cytometry was used to analyze PBMCs from study participants for effector and memory T cell populations, including CD4^+^ and CD8^+^ central memory T cells (T_CM_), Naïve T cells, effector memory T cells (T_EM_), and effector memory CD45RA^+^ T cells (T_EMRA_) immune cell markers.

**Results:** CD8^+^ T_EMRAs_ were positively correlated with Nf-L and GFAP. We found no significant difference in CD8^+^ T_EMRAs_ based on cognitive scores and no associations between CD8^+^ T_EMRAs_ and AD-related biomarkers. CD4^+^ T_EMRAs_ were associated with cognitive impairment on the MMSE. Gender was not associated with T_EMRAs_, but it did show an association with other T cell populations.

**Conclusion:** These findings suggest that the accumulation of CD8^+^ T_EMRAs_ may be a response to neuronal injury (Nf-L) and neuroinflammation (GFAP) during aging or the progression of AD and ADRD. As our findings in a community-based cohort were not clinically-defined AD participants but included all ADRDs, this suggests that T_EMRAs_ may be associated with changes in systemic immune T cell subsets associated with the onset of pathology.

## Introduction

The innate and adaptive immune system, essential for protecting individuals against infections, injury, and diseases, is profoundly impacted by chronological aging, becoming less protective and more inflammatory[1]. The age-associated decline in T cell function contributes to increased susceptibility to infections such as influenza, COVID-19, pneumonia, and urinary tract infections (UTI) [1, 2]. Beyond aging, changes in adaptive immune responses are exacerbated in Alzheimer’s disease and related dementias (ADRD) [3–10]. The ADRD-associated adaptive immune changes include a decrease in naïve T cells and the accumulation of cytotoxic memory and effector memory T cells [3–9], which may further weaken the protection against pathogens and vaccine responsiveness. Infections such as pneumonia and UTI are higher in individuals with dementia[11], with the weakened immune system being a major contributor to the greater infection-induced mortality in individuals with dementia [12, 13].

A handful of studies have evaluated adaptive immune response in ADRD, highlighting a central role for terminally differentiated effector memory T cells (called T_EMRAs_). T_EMRAs_ are CD4^+^ or CD8^+^ T cells that re-express CD45RA. In clinic-based studies, more T_EMRAs_ were associated with increased neurodegenerative pathology and worse cognitive function [5–7].

TEMRAs, which are also associated with viral infections such as cytomegalovirus (CMV), dengue, and human immunodeficiency virus (HIV) [3, 14, 15], are known for their T cell effector function, low proliferation capacity, and cytotoxic activity [16–18].

While changes in T cell subsets, including T_EMRAs_, are linked to Alzheimer’s disease (AD) [5–7], studies have not made it clear if these changes are specific to AD-related cognitive decline or are more generally correlated with dementia, as prior studies were limited to people with AD-related dementia diagnosed by reduced cognitive scores, computed tomography (CT), magnetic resonance scanning (MRI) for volumetric loss of brain regions, Aβ PET imaging, and reduced CSF Aβ and tau levels[5–7]. In addition, the connection between T_EMRAs_ and early stages of cognitive decline is not well established. We analyzed cells by flow cytometry from two groups, cognitively healthy control participants (clinical dementia rating (CDR) = 0) and individuals with early stages of cognitive decline (CDR =0.5-1), to identify associations between CD4^+^ or CD8^+^ T_EMRAs_ in peripheral blood and early cognitive decline in a community-based cohort of older individuals. To test associations between T_EMRAs_ and AD, we measured plasma biomarkers of AD (Aβ & tau), neuronal injury (Nf-L), and astrocyte-mediated neuroinflammation (GFAP). We found that T_EMRAs_ are associated with low MMSE, biomarkers of AD (Aβ 42/40), and brain injury (Nf-L and GFAP).

## Materials and Methods

### Human subjects

The participants in this study were recruited from a larger cohort study of aging and dementia at the University of Kentucky Alzheimer’s Disease Research Center [19]. Before participating in the study, the participants provided informed consent according to University of Kentucky Institutional Review Board-approved protocols. The AD Research Center recruits older adults from central Kentucky and includes approximately 500 active participants who receive annual follow-up visits. These visits include a variety of measures, such as physical and neurological examinations, medical histories, cognitive testing, and fluid biomarkers. For our analysis, we recruited n = 40 cognitively healthy individuals (CDR = 0) and n = 44 cognitively impaired (CDR=0.5-1) individuals, with similar numbers based on gender, and cardiovascular disease risk factors and age. The participants were excluded if they had type 2 diabetes, auto-immune disease, or active cancer. Blood samples were collected from 84 participants between August 2020 and June 2021, and clinical evaluations and cognitive testing were performed using the National Alzheimer’s Coordinating Center uniform data set [20, 21]. The de-identified samples were provided to the experimentalists, who were blinded to the participant group identity until all data had been generated.

### Peripheral blood mononuclear cell (PBMC) isolation

5-10mls of peripheral blood was taken through venous puncture and stored in acid/citrate/dextrose tubes. PBMCs were isolated by density centrifugation in Ficoll histopaque 1077 using SepMate PBMC isolation tube (StemCell tech, Cat#: 85415) according to the manufacturer’s protocol. After the cells were washed in buffer (0.1 BSA, 2mM EDTA, 1X PBS), residual red blood cells were removed using red blood cells lysis buffer (EBioscience, cat#: 00- 4300-54). The cells were then frozen at a density of 1x10^6^/ml in freezing media containing (90%FBS Heat Inactivated (HI), 10% DMSO) and stored at -80°C in a Mr. Frosty apparatus (Nalgene) for 24 hours before being transferred to long-term storage in liquid N2 until use as previously described [22].

### Plasma biomarker analysis

The plasma collected from the same blood drawn as the PBMCs underwent analysis on the Quanterix Simoa HD-X for plasma biomarkers as previously described [23].

### Flow cytometry

After being thawed, PBMCs were washed with warm RPMI followed by PBS. The cells were then stained with viability marker Ghost dye-Alexa Fluor 700 (Tonbo Biosciences) for 30 minutes at 4 °C, then washed with FACS buffer (1X PBS supplemented with 1% bovine serum albumin and .1% sodium azide). To prevent unintended antibody binding to Fc receptors, cells were blocked with FcR blocking reagent, human (Miltenyi Biotec) for 10 minutes at 4 °C. The cells were then stained at a 1:50 dilution with antibodies to immune cell targets as follows: CD3- FITC (BD Biosciences, 555332), CD8-PE (BD Biosciences, 555635), CD4-APC (BD Biosciences, 555349), CCR7-buv395 (BD Biosciences, 568681), CD45RA-bv421 (BD Biosciences, 569620) for 30 minutes at 4 °C. The cells were then washed twice with FACS buffer and fixed using cytofix from BD Biosciences for 30 minutes at 4 °C, followed by a wash with FACS buffer and resuspension in FACS buffer. Cell populations were acquired using the BD Symphony A3, and Flow v10.8 software was used for analysis. Fluorescence-minus-one controls were utilized to identify cell populations, and each experiment had a single-stained control and negative control.

### Statistics

Characteristics of participants were assessed with analysis of variance or χ2 tests. Fluid biomarker data were log2 transformed. For categorical comparisons, a parametric t-test or one-way ANOVA were used for the combined data, or the data stratified by gender. Multiple linear regression models were used to adjust mean responses for covariates, as listed in figures and tables. Pairwise effect sizes were calculated using Cohen’s d. The effect size for linear regressions were calculated as R^2^ or **W_2_.** For the box-and-whiskers graph, the box shows the median, 25th and 75th percentile, and the whiskers show the minimum and maximum values. Data visualizations were created in JMP Pro v16 or GraphPad Prism v9.4.

## Results

### Patient Characteristics

The study included 84 participants from the UK-ADRC community cohort, comprising 40 cognitively healthy (mean age 77, range 62-95) and 44 cognitively impaired (mean age 79, range 64-97) individuals, balancing for gender, cognitive status (CDR=0 vs. CDR=0.5-1), age, and cardiovascular risk factors, with demographic details in Table 1 and no marked differences by CDR between groups.

**Table 1.** Demographic information.

|  | CDR = 0 | CDR = 0.5-1 |
| --- | --- | --- |
| Number of participants, (n) | 40 | 44 |
| Age in years (mean ± SD) | 77±7.9 | 79±7.9 |
| Gender (Men: Women) | 16:24 | 24:21 |
| APOE ε4 carrier (n) | 9 (22.5%) | 20 (48.9%) |
| Hypertension (n) | 24 (60%) | 25 (55.5%) |
| Hyperlipidemia (n) | 25 (62.5%) | 27 (60%) |
| BMI (mean ± SD) | 26.4±3.5 | 25.7±4.3 |
| Education (mean ± SD) | 17±2.5 | 16.7±2.9 |
| MMSE (mean ± SD) | 29±1.2 | 23.9±5.9 |
| MoCA (mean ± SD) | 26.8±2.2 | 18.5±6.2 |
| CD3<sup>+</sup> (%) (mean ± SD) | 72.3 ± 9.9 | 68.8 ± 12.2 |
| CD3<sup>+</sup> CD4<sup>+</sup> (%) (mean ± SD) | 73.5 ± 13.2 | 71.0 ± 13.1 |
| CD3<sup>+</sup> CD8<sup>+</sup> (%) (mean ± SD) | 22.8 ± 11.3 | 25.6 ± 12.4 |
| CD4<sup>+</sup>/CD8<sup>+</sup> (mean ± SD) | 4.9 ± 4.1 | 4.0 ± 3.6 |

Abbreviations: Clinical Dementia Rating (CDR), body mass index (BMI), Mini-Mental State Examination (MMSE), Montreal Cognitive Assessment (MoCA)

### Higher Percentage of CD4^+^ T_EMRAs_ in Individuals with low MMSE

Prior studies showed an association between greater CD8^+^ T_EMRAs_ and worse cognitive function in people with probable AD, and greater CD8^+^ T_EMRAs_ in cognitively healthy people with preclinical AD, defined by cognitive scores, CSF Aβ and tau levels, MRI, and Aβ PET imaging [5, 6]. Yet, it is unclear if changes in CD8^+^ T_EMRAs_ are an AD-specific phenotype or are seen more generally with cognitive decline, and if the changes are limited to CD8^+^ T cells or are also seen in CD4^+^ T cells. As T_EMRAs_ are seen even in preclinical AD, we also wanted to test in a community-based cohort if T_EMRAs_ are seen with early changes in global cognitive function.

Using flow cytometry, we analyzed CD3^+^CD4^+^ and CD3^+^CD8^+^ T cells **(Figure 1A**, **Table 1)** for naïve and memory phenotypes based on CD45RA and CCR7 staining **(Figure 1A)**. CD4^+^ and CD8^+^ T_EMRAs_ were not associated with CDR **(Figure 1B)**, which held when adjusted for age and gender (Table 2). CD4^+^ T_EMRAs_ (p=0.0292) and CD8^+^ T_EMRAs_ (p=0.0779) were negatively associated with MMSE scores <25 **(Figure 1C)**. When adjusted for age and gender, significance was lost between MMSE and T_EMRAs_; however, the trend for a change in T_EMRAs_ by MMSE held for both CD4^+^ T_EMRAs_ (p=0.0531) and CD8^+^ T_EMRAs_ (p=0.101) **(Table 2)**. There was no association between CD4^+^ T_EMRAs_ and CD8^+^ T_EMRAs_ and MoCA **(Figure 1D**, **Table 2)**. These results suggest that in a community-based cohort, the increased frequency of T_EMRAs_ occurs with greater severity of cognitive decline and may be associated with AD-related cognitive decline, which only accounts for a proportion of the cognitive decline in our cohort, thus diluting the effect size seen in our study.

**Figure 1:**
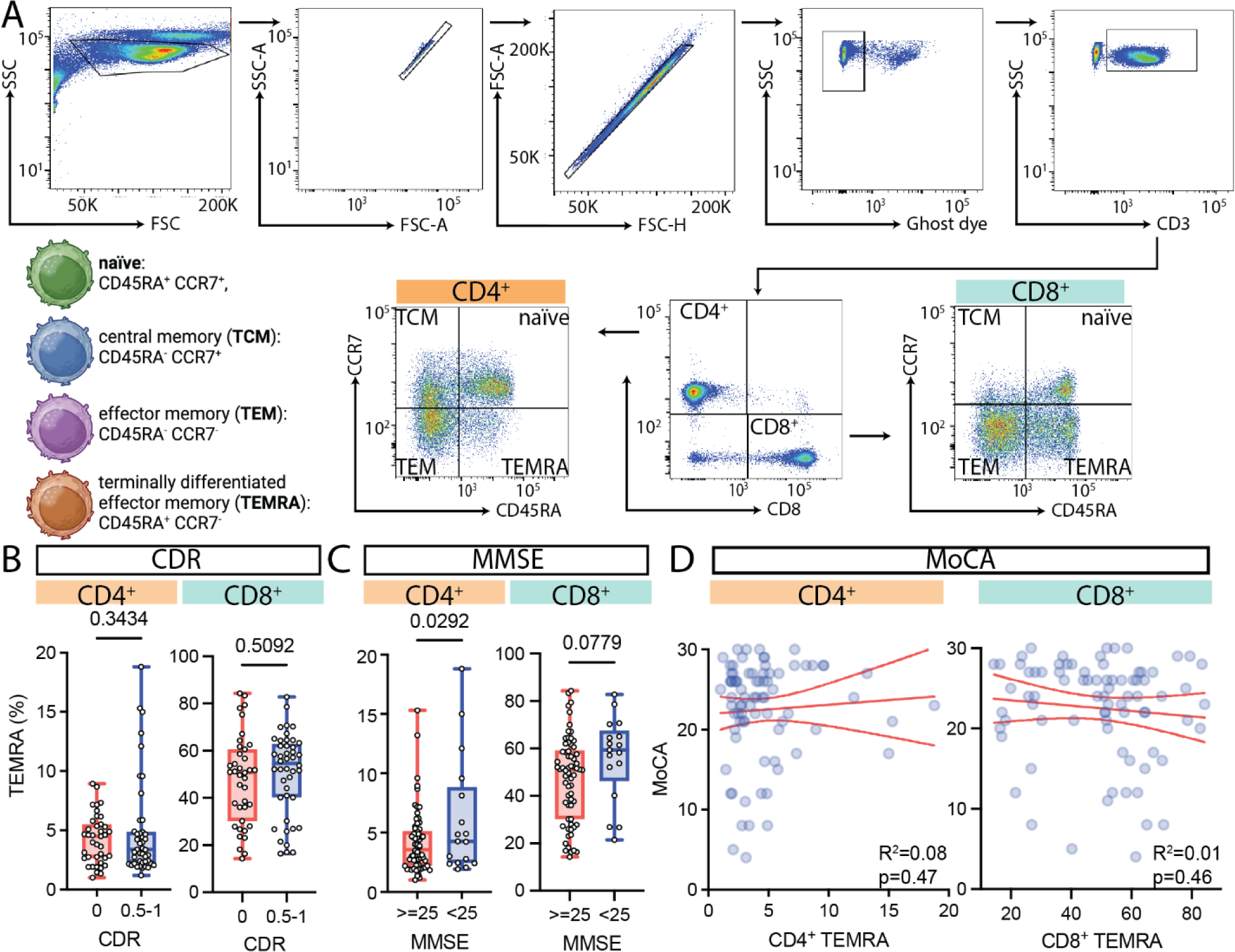
Association of CD4^+^ and CD8^+^ T_EMRAs_ with global cognitive function. (A) Representative flow cytometry gating strategy for lymphocyte subsets. The strategy begins with gating on lymphocytes, excluding doublets and dead cells. T cells are gated on CD3^+^; CD4^+^ T cells are CD3^+^ CD4^+^; CD8^+^ T cells are CD3^+^ CD8^+^. The CD4^+^ and CD8^+^ T cells are then further divided into central memory (T_CM_): CD45RA^-^CCR7^+^, naive: CD45RA^+^CCR7^+^, effector memory (T_EM_): CD45RA^-^CCR7^-^, and terminally differentiated effector memory re-expressing CD45RA (T_EMRA_): CD45RA^+^CCR7^-^. Association of CD4^+^ and CD8^+^ TEMRAs with **(B)** CDR **(C)** MMSE and **(D)** MoCA. Circles represent individual participants. The dashed line shows the 95% confidence interval for the linear regression. See also, Table 2.

**Table 2:** Association between measures of global cognitive function and T cell subsets.

| T cell subset |  | TCM |  | Naïve |  | TEMRA |  | TEM |  |
| --- | --- | --- | --- | --- | --- | --- | --- | --- | --- |
| Model |  | 1 | 2 | 1 | 2 | 1 | 2 | 1 | 2 |
| **CDR**<br>(p-value;<br>Cohens D) | **CD4<sup>+</sup>** | 0.254;<br>0.253 | 0.45;<br>0.169 | 0.369;<br>0.199 | 0.573;<br>0.126 | 0.343;<br>0.21 | 0.263;<br>0.251 | 0.979;<br>0.006 | 0.863;<br>0.038 |
|  | **CD8<sup>+</sup>** | 0.368;<br>0.199 | 0.314;<br>0.226 | 0.407;<br>0.183 | 0.741;<br>0.074 | 0.509;<br>0.146 | 0.641;<br>0.104 | 0.708;<br>0.083 | 0.941;<br>0.016 |
| **MMSE**<br>(p-value;<br>Cohens D) | **CD4<sup>+</sup>** | **0.009**;<br>0.725 | **0.013**;<br>0.702 | 0.101;<br>0.44 | 0.135;<br>0.415 | **0.029**;<br>0.604 | 0.053;<br>0.54 | 0.465;<br>0.2 | 0.529;<br>0.174 |
|  | **CD8<sup>+</sup>** | 0.281;<br>0.295 | 0.331;<br>0.269 | 0.412;<br>0.224 | 0.339;<br>0.264 | 0.078;<br>0.486 | 0.101;<br>0.456 | 0.244;<br>0.319 | 0.351;<br>0.258 |
| **MoCA**<br>(p-value;<br>R<sup>2</sup>) | **CD4<sup>+</sup>** | 0.664;<br>0.05 | 0.59;<br>0.09 | 0.58;<br>0 | 0.52;<br>0.06 | 0.47;<br>0.08 | 0.51;<br>0.09 | 0.51;<br>0.01 | 0.48;<br>0.04 |
|  | **CD8<sup>+</sup>** | 0.36;<br>0.02 | 0.35;<br>0.02 | 0.64;<br>0 | 0.83;<br>0.11 | 0.46;<br>0.01 | 0.6;<br>0.04 | 0.85;<br>0 | 0.84;<br>0.06 |

Model 1 is unadjusted. Model 2 includes gender and age. Abbreviations: Clinical Dementia Rating (CDR), Mini-Mental State Examination (MMSE), Montreal Cognitive Assessment (MoCA)

### Associations of T_EMRAs_ with other relevant biological variables

To determine the impact of other biological variables, including gender, age, body mass index (BMI), and apoE genotype, on the frequency of CD4^+^ T_EMRAs_ and CD8^+^ T_EMRAs_, we completed regression models. We found no associations between apoE or BMI and T cell phenotypes **(Table 3)**. Similarly, age was not associated with CD4^+^ T_EMRAs_ and CD8^+^ T_EMRAs_ **(Table 3)**. However, age negatively correlated with naïve CD8^+^ T cells (p=0.007) **(Table 3)**. While there were no differences in the frequencies of CD4^+^ T_EMRAs_ and CD8^+^ T_EMRAs_ by gender, we detected a higher frequency of CD4^+^ (p=0.0369) and CD8^+^ (p=0.0192) naïve T cells in women while men have a higher frequency of CD4^+^ T_CMs_ (0.0056) and CD8^+^ T_EMs_ (p=0.0137) **(Figure 2)**.

**Figure 2:**
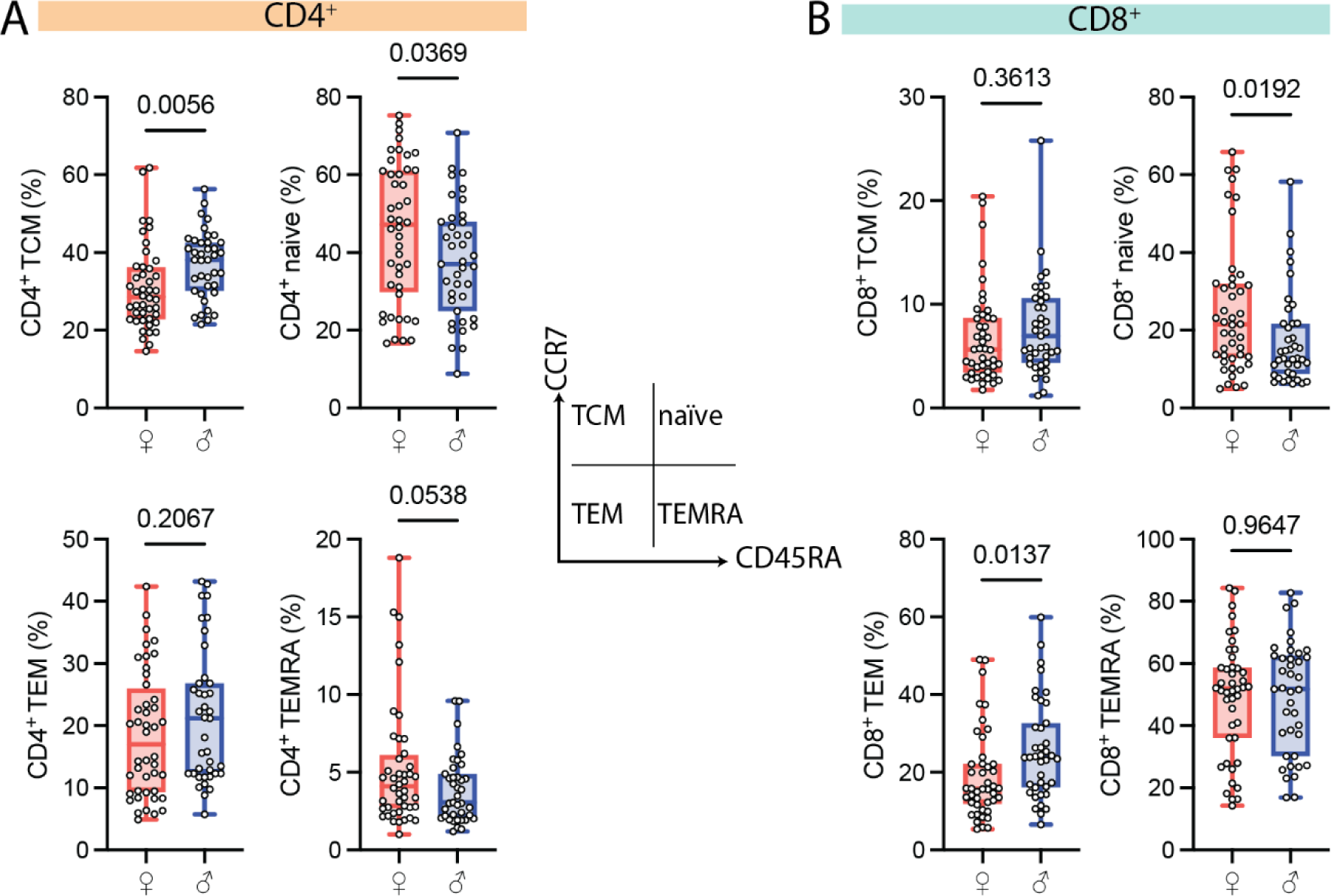
Association of gender with T cell subsets. The frequency of CD4^+^ **(A)** and CD8^+^ **(B)** T-cell subsets in the blood of women (♀) and men (♂). Circles represent individual participants. See also, Table 3.

**Table 3:** Association between other biological variables and T cell subsets.

| T cell subset |  | TCM |  | Naïve |  | TEMRA |  | TEM |  |
| --- | --- | --- | --- | --- | --- | --- | --- | --- | --- |
| Model |  | 1 | 2 | 1 | 2 | 1 | 2 | 1 | 2 |
| **Gender**<br>(p-value;<br>Cohens D) | **CD4<sup>+</sup>** | **0.006;**<br>0.626 | **0.01;**<br>0.586 | **0.037;**<br>0.467 | 0.06;<br>0.426 | 0.054;<br>0.43 | **0.045;**<br>0.454 | 0.21;<br>0.28 | 0.246;<br>0.261 |
|  | **CD8<sup>+</sup>** | 0.36;<br>0.202 | 0.37;<br>0.201 | **0.019;**<br>0.525 | **0.047;**<br>0.449 | 0.965;<br>0.01 | 0.803;<br>0.056 | **0.014;**<br>0.554 | **0.017;**<br>0.54 |
| **Age**<br>(p-value;<br>R<sup>2</sup>) | **CD4<sup>+</sup>** | 0.139;<br>0.027 | 0.284;<br>0.104 | 0.165;<br>0.024 | 0.285;<br>0.066 | 0.76;<br>0.001 | 0.51;<br>0.05 | 0.499;<br>0.005 | 0.639;<br>0.022 |
|  | **CD8<sup>+</sup>** | 0.89;<br>0 | 0.99;<br>0.01 | **0.007;**<br>0.085 | **0.018;**<br>0.13 | 0.09;<br>0.035 | 0.09;<br>0.04 | 0.5;<br>0.006 | 0.78;<br>0.073 |
| **BMI**<br>(p-value;<br>R<sup>2</sup>) | **CD4<sup>+</sup>** | 0.284;<br>0.014 | 0.575;<br>0.107 | 0.983;<br>0 | 0.605;<br>0.07 | 0.55;<br>0 | 0.61;<br>0.05 | 0.37;<br>0.01 | 0.23;<br>0.04 |
|  | **CD8<sup>+</sup>** | 0.425;<br>0.007 | 0.48;<br>0.02 | 0.41;<br>0.008 | 0.08;<br>0.16 | 0.21;<br>0.02 | 0.09;<br>0.07 | 0.6;<br>0.003 | 0.85;<br>0.07 |
| **ApoE**<br>(p-value;<br>R<sup>2</sup>) | **CD4<sup>+</sup>** | 0.222;<br>0.304 | 0.411;<br>0.207 | 0.57;<br>0.141 | 0.836;<br>0.052 | 0.103;<br>0.407 | 0.16;<br>0.354 | 0.85;<br>0.047 | 0.97;<br>0.008 |
|  | **CD8<sup>+</sup>** | 0.237;<br>0.294 | 0.3;<br>0.262 | 0.413;<br>0.203 | 0.73;<br>0.086 | 0.79;<br>0.067 | 0.68;<br>0.104 | 0.31;<br>0.25 | 0.51;<br>0.167 |

Model 1 is unadjusted. Model 2: For gender includes age. For age includes gender. For BMI and ApoE includes age and gender. ApoE is the presence of 1 or more *E4* alleles. *E4* = 29 participants, non-*E4* = 40 participants, ApoE data unavailable for 15 participants.

### T_EMRAs_ correlated with plasma markers of neurodegeneration and neuroinflammation

Prior studies stratified their study participants as either controls, or clinical / biomarker defined preclinical AD or probable AD [5–7]. Our community-based study, was stratified by cognitive status but was not limited to AD. Therefore, we tested if plasma biomarkers AD (Aβ42/Aβ40 ratio, and total-tau) and of neurodegeneration and brain injury (Nf-L and GFAP), would show a greater association with T_EMRAs_ then what was seen for cognitive status.

Aβ42/Aβ40 ratio was positively correlated with the frequency of CD4^+^ T_EMRAs_ (p=0.036, **Figure 3A**); however, this association did not hold following adjustments for age and gender **(Table 4)**. We found no association between the frequency of CD8^+^ T_EMRAs_ and the Aβ42/Aβ40 ratio **(Figure 3B**, **Table 4).** CD8^+^ naïve T cells correlated negatively with Aβ42/Aβ40 ratio; however, this correlation was not significant after accounting for age and gender **(Table 4)**. While there was no association between CD4^+^ T_EMRAs_ **(Figure 3C**, **Table 4)** and CD8^+^ T_EMRAs_ with total tau **(Figure 3D**, **Table 4)**, we identified a negative correlation between total tau and CD4^+^ central memory T cells and a positive correlation between total tau and CD4^+^ naïve T cells **(Table 4)**.

**Figure 3:**
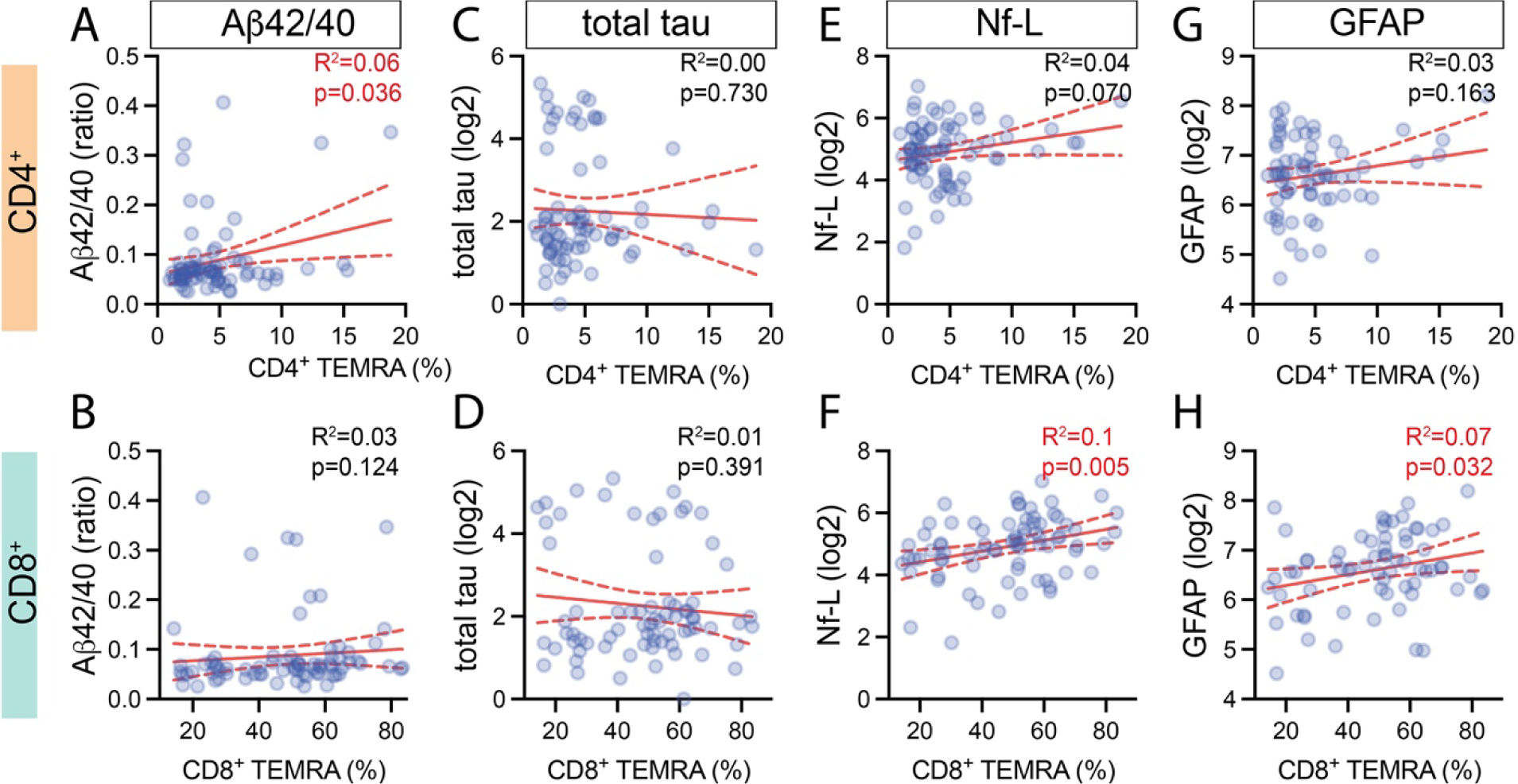
T_EMRAs_ correlated with plasma markers of neurodegeneration and neuroinflammation. Correlation between the percentage of CD4^+^ and CD8^+^ TEMRA s in the blood of individuals and plasma levels of AD-related biomarkers (A,B) Aβ42/40, (C,D) total tau, (E,F) Nf-L, and (G,H) GFAP. The dashed line shows the 95% confidence interval for the linear regression. The p-values and R^2^ values shown are for the unadjusted analysis. See also Table 4. Markers are for individual participants.

**Table 4:** Association between other biological variables and T cell subsets.

| T cell subset |  | TCM |  | Naïve |  | TEMRA |  | TEM |  |
| --- | --- | --- | --- | --- | --- | --- | --- | --- | --- |
| Model |  | 1 | 2 | 1 | 2 | 1 | 2 | 1 | 2 |
| **Aβ42/40**<br>(p-value; R<sup>2</sup>) | **CD4<sup>+</sup>** | 0.955;<br>0 | 0.785;<br>0.117 | 0.221;<br>0.02 | 0.306;<br>0.074 | **0.02**;<br>0.072 | **0.032**;<br>0.131 | 0.193;<br>0.023 | 0.221;<br>0.033 |
|  | **CD8<sup>+</sup>** | 0.938;<br>0 | 0.91;<br>0.003 | 0.096;<br>0.037 | 0.175;<br>0.168 | 0.451;<br>0.008 | 0.681;<br>0.04 | 0.357;<br>0.012 | 0.343;<br>0.073 |
| **total tau**<br>(p-value; R<sup>2</sup>) | **CD4<sup>+</sup>** | **0.036**;<br>0.06 | 0.08;<br>0.153 | **0.023**;<br>0.07 | 0.058;<br>0.108 | 0.73;<br>0.002 | 0.87;<br>0.066 | 0.145;<br>0.029 | 0.199;<br>0.035 |
|  | **CD8<sup>+</sup>** | 0.681;<br>0.002 | 0.672;<br>0.005 | 0.052;<br>0.051 | 0.207;<br>0.166 | 0.392;<br>0.01 | 0.711;<br>0.037 | 0.186;<br>0.024 | 0.261;<br>0.073 |
| **Nf-L**<br>(p-value; R<sup>2</sup>) | **CD4<sup>+</sup>** | 0.619;<br>0.003 | 0.814;<br>0.086 | **0.05**;<br>0.051 | 0.058;<br>0.088 | 0.07;<br>0.044 | 0.11;<br>0.086 | **0.037**;<br>0.057 | **0.022**;<br>0.078 |
|  | **CD8<sup>+</sup>** | 0.928;<br>0 | 0.908;<br>0.015 | **0.006**;<br>0.098 | **0.03**;<br>0.181 | **0.005**;<br>0.102 | **0.021**;<br>0.104 | 0.506;<br>0.006 | 0.423;<br>0.071 |
| **GFAP**<br>(p-value; R<sup>2</sup>) | **CD4<sup>+</sup>** | 0.524;<br>0.006 | 0.295;<br>0.153 | 0.583;<br>0.005 | 0.804;<br>0.055 | 0.163;<br>0.029 | 0.384;<br>0.084 | 0.315;<br>0.015 | 0.301;<br>0.022 |
|  | **CD8<sup>+</sup>** | 0.497;<br>0.007 | 0.582;<br>0.029 | 0.053;<br>0.055 | 0.149;<br>0.144 | **0.032**;<br>0.068 | 0.086;<br>0.074 | 0.598;<br>0.004 | 0.536;<br>0.042 |

Model 1 is unadjusted. Model 2 includes age and gender. Sample size of analysis follows: Aβ42/Aβ40 (n=75), total tau (n=75), Nf-L (n=77), GFAP (n=69). Abbreviations: neurofilament light (Nf-L), glial fibrillary acidic protein (GFAP).

These results suggest that the change in T_EMRAs_ may be more indicative of neurodegeneration in general, than an AD-specific change.

Next, we tested the association of T cell subsets and brain injury biomarkers Nf-L **(Figure 3E,F)** and GFAP **(Figure 3G,H)**. Frequency of CD8^+^ T_EMRAs_ **(p=0.005; Figure 3F)** but not CD4^+^ T_EMRAs_ **(p=0.07 Figure 3E)** positively correlated with plasma Nf-L. Plasma Nf-L positively correlated with CD4^+^ central memory T cells **(Table 4)**, however, we found negative correlations between Nf-L and CD4^+^ and CD8^+^ naïve T cells **(Table 4)**. Plasma GFAP levels correlated with CD8^+^ T_EMRAs_ **(p=0.032 Figure 3H)**, but not CD4^+^ T_EMRAs_ **(p=0.163 Figure 3G)**. We conclude that CD8^+^ T_EMRAs_ are associated with early neurodegeneration and/or acute brain injury in the study participants.

## Discussion

Terminally differentiated effector memory T cells have been linked to AD-related cognitive decline[5–7]. This study aimed to test whether worse cognitive function was associated with greater proportions of these T_EMRAs_ in a community-based cohort of older individuals. We found more CD4^+^ T_EMRAs_ in individuals with lower MMSE scores. We also found that T cell subsets varied by gender, but not specifically for CD4^+^ T_EMRAs_ and CD8^+^ T_EMRAs_. In our study, CD8^+^ T_EMRAs_, previously linked to AD-related cognitive decline defined by CSF AD-biomarkers, cognition, and imaging [5–7], correlate with plasma biomarkers of brain injury (Nf-L and GFAP). In our community-based aging cohort, we also found that terminally differentiated effector memory T cells were also associated with both cognitive impairment and brain injury biomarkers. Our findings suggest that T_EMRAs_ may be related to changes in systemic immune T cell subsets associated with neuronal injury in dementia, including AD.

Studies from multiple groups have shown that CD8^+^ T_EMRAs_ accumulate in the blood of pre-clinical, MCI, and AD patients[5–7]. Although we did not see significant increases in CD8^+^ T_EMRAs_ in our cognitively impaired participants classified by CDR and MoCA, we note a trend towards more CD8^+^ T_EMRAs_ in our participants with low MMSE scores. The lack of difference could be because the cognitive decline seen in our participants could be from multiple etiologies. Prior studies only included patients positive for Aβ, probable AD, or MCI due to AD [5–7], which suggests that the accumulation of CD8^+^ T_EMRAs_ seen may be specific to AD. However, increases in CD8^+^ T_EMRAs_ were also seen in cognitively healthy individuals with PET imaging positive for Aβ[6], implying that an individual doesn’t have to be cognitively impaired to have a higher frequency of CD8^+^ T_EMRAs_. Thus, the accumulation of CD8^+^ T_EMRAs_ may be more associated with the pathology or injury caused by the accumulation of Aβ than overt cognitive decline.

Our lack of differences in CD8^+^ T_EMRAs_ led us to examine the influence of other important biological variables. In contrast to other studies, we did not find any significant differences in CD8^+^ T_EMRAs_ by gender, however in line with previous studies, women in the cohort, irrespective of cognitive status, have significantly more CD4^+^ and CD8^+^ naïve T cells[3]. Women in the cohort also have significantly fewer CD4^+^ central memory and CD8^+^ effector memory T cells compared to men. The difference in naïve populations suggests that most women in our cohort may have more T cells that are not cytotoxic and can respond to new antigens; however, this was not tested. More effector memory T cells in men suggest that they are in a more cytotoxic state, with populations that can secrete proinflammatory cytokines or are ready for antigen restimulation. Overall, however, the increased percentage of CD8^+^ naïve T cells potentially contributes to our lack of differences in our CD8^+^ T_EMRAs_.

CD4^+^ T_EMRAs_ in healthy individuals are found in the blood at a lower frequency (0.3% to 18%[24]) than their CD8^+^ counterparts. The two cell types share similarities, including the ability to secrete perforin and granzyme and relatively reduced proliferative capacities[16]. In contrast to the CD8^+^ T_EMRAs_, we did see more CD4^+^ T_EMRAs_ in participants with lower MMSE scores, which agrees with previous studies showing more CD4^+^ T_EMRAs_ in people with AD [7]. CD4^+^ memory and cytotoxic T cells have also been shown to be associated with patients with Parkinson’s disease [8–10], suggesting a degree of specificity in T cell changes to different neurodegenerative diseases. Our results agree with this interpretation as the early stages of cognitive decline, and the multiple possible disease etiologies causing that decline in our study participants could lead to mixed CD4^+^ and CD8^+^ responses in our group analysis. Future studies using pathologically-confirmed consensus diagnosis would test this hypothesis. Recent studies also show a role for mitochondria in T_EMRAs_, with CD4^+^ T_EMRAs_ having more mitochondrial content and fewer mitochondrial nodes when compared to CD8^+^ T_EMRAs_[25]. As different mitochondria abnormalities are seen in PD vs AD, for example, these changes in mitochondria could be responsible for potential specificity in T cell changes to different neurodegenerative diseases.

While studies have shown evidence of TEMRA-induced protective functions in the setting of dengue[14, 16], the function of these cells in neurodegenerative disease is unknown. Previous studies found evidence of CD8^+^ T cells in the brain with AD pathology and suggested that these cells contribute to neuronal injury and neuroinflammation [5, 26]. In agreement with this suggestion, we found a positive correlation between CD8^+^ T_EMRAs_ and plasma biomarkers of neuronal injury (Nf-L) and neuroinflammation (GFAP). In a mouse model of tauopathy, depleting T cells or using an immune checkpoint blockade (PDCD1) reduced neurodegeneration [26], which agrees with other studies in models of neurodegenerative disease[27–29], suggesting a provocative hypothesis that T cells could be directly causing neurodegeneration in AD.

In contrast to a protective function, increased abundance of circulating T_EMRAs_ is also associated with high cytotoxicity and anti-viral functions[16–18]. When CD4^+^ and CD8^+^ T cells differentiate into T_EMRAs_, they stop expressing the co-stimulatory molecules CD28 and CD27,molecules crucial for T-cell activation, proliferation, and survival[30]. T_EMRAs_ upregulate CD57, a senescence marker associated with decreased telomere length and low proliferative capacity[31]. Viral infections lead to the accumulation of CD8^+^ T_EMRAs_[32, 33], leading to the idea that the accumulation of T_EMRAs_ during aging and the progression of AD could be due to latent viral infections[3, 5, 6]. While studies have shown that infections with viruses such as herpes simplex virus (HSV) and CMV are associated with the development of AD[34, 35], recent studies in pre-clinical AD find the accumulation of CD8^+^ T_EMRAs_ was independent of latent, reactivated, or acute Herpesviridae background infections[6]. This suggests that T_EMRAs_ can also accumulate in the absence of viral infections, again highlighting a potential neuroinjury response.

Our study has limitations. Firstly, our participants are from a community-based cohort at a single site. While we tried to balance cardiovascular disease risk factors between the groups, one study can never entirely represent the community cohort. Moreover, the cohort mainly comprises highly educated white individuals, which does not reflect the broader elderly population. Future studies must be done in a larger cohort with more racial, ethnic, and socioeconomic diversity. Furthermore, we did not assay for any viral infection, so we don’t know if any participants in our group had latent or prior viral infections such as CMV, HIV, or HSV, which could skew TEMRA populations in both cognitively normal and impaired people.

In summary, the accumulation of CD8^+^ T_EMRAs_ is either specific for AD, as shown by other[5–7], or is associated with neuronal injury and neuroinflammation caused by ADRD. CD4^+^ T_EMRAs_ may also be important in progressing AD and AD-related dementia. As we see more CD4^+^ T_EMRAs_ in our cohort, future studies should focus on whether CD4^+^ T_EMRAs_ are an early sign of AD or ADRD progression. Furthermore, since CD4^+^ T cells are well known to assist with CD8^+^ T cell memory function, studies should determine how the presence of CD4^+^ T_EMRAs_ impacts the accumulation of CD8^+^ T_EMRAs_ to determine the interplay between the two. The question remains if their relationship is dependent or independent of each other, and how it contributes to ongoing pathology, though, a better understanding of these critical immune populations could shed light on potential therapeutic strategies for millions at risk for ADRD.

### Data availability

All data generated in this study are available in the main text. Additional data are available following reasonable request to the University of Kentucky Alzheimer Disease Research Center.

## Acknowledgements

Our sincere thanks go out to all the research participants and their families. Illustrations were created using BioRender.com. We would like to thank Tiffany L. Sudduth with her help with the plasma biomarker data.

## References

1. Lutshumba, J., B.S. Nikolajczyk, and A.D. Bachstetter, Dysregulation of Systemic Immunity in Aging and Dementia. Front Cell Neurosci, 2021. 15: p. 652111.

2. Guo, L., et al., T cell aging and Alzheimer’s disease. Front Immunol, 2023. 14: p. 1154699.

3. Fang, Y., et al., Circulating immune cell phenotypes are associated with age, sex, CMV, and smoking status in the Framingham Heart Study offspring participants. Aging (Albany NY), 2023. 15(10): p. 3939–3966.

4. Thyagarajan, B., et al., Age-Related Differences in T-Cell Subsets in a Nationally Representative Sample of People Older Than Age 55: Findings From the Health and Retirement Study. J Gerontol A Biol Sci Med Sci, 2022. 77(5): p. 927–933.

5. Gate, D., et al., Clonally expanded CD8 T cells patrol the cerebrospinal fluid in Alzheimer’s disease. Nature, 2020. 577(7790): p. 399-404.

6. Gericke, C., et al., Early beta-amyloid accumulation in the brain is associated with peripheral T cell alterations. Alzheimers Dement, 2023.

7. Larbi, A., et al., Dramatic shifts in circulating CD4 but not CD8 T cell subsets in mild Alzheimer’s disease. J Alzheimers Dis, 2009. 17(1): p. 91–103.

8. Saunders, J.A., et al., CD4+ regulatory and effector/memory T cell subsets profile motor dysfunction in Parkinson’s disease. J Neuroimmune Pharmacol, 2012. 7(4): p. 927–38.

9. Wang, P., et al., Single-cell transcriptome and TCR profiling reveal activated and expanded T cell populations in Parkinson’s disease. Cell Discov, 2021. 7(1): p. 52.

10. Contaldi, E., L. Magistrelli, and C. Comi, T Lymphocytes in Parkinson’s Disease. J Parkinsons Dis, 2022. 12(s1): p. S65–S74.

11. Ryvicker, M., et al., Clinical and Demographic Profiles of Home Care Patients With Alzheimer’s Disease and Related Dementias: Implications for Information Transfer Across Care Settings. J Appl Gerontol, 2022. 41(2): p. 534–544.

12. Pereira, B., X.N. Xu, and A.N. Akbar, Targeting Inflammation and Immunosenescence to Improve Vaccine Responses in the Elderly. Front Immunol, 2020. 11: p. 583019.

13. Gilstrap, L., et al., Trends in Mortality Rates Among Medicare Enrollees With Alzheimer Disease and Related Dementias Before and During the Early Phase of the COVID-19 Pandemic. JAMA Neurol, 2022. 79(4): p. 342–348.

14. Tian, Y., et al., Unique phenotypes and clonal expansions of human CD4 effector memory T cells re-expressing CD45RA. Nat Commun, 2017. 8(1): p. 1473.

15. Meraviglia, S., et al., *T-Cell Subsets (T(CM), T(EM),* T(EMRA)) and Poly-Functional Immune Response in Patients with Human Immunodeficiency Virus (HIV) Infection and Different T-CD4 Cell Response. Ann Clin Lab Sci, 2019. 49(4): p. 519–528.

16. Patil, V.S., et al., Precursors of human CD4(+) cytotoxic T lymphocytes identified by single-cell transcriptome analysis. Sci Immunol, 2018. 3(19).

17. Appay, V., et al., Phenotype and function of human T lymphocyte subsets: consensus and issues. Cytometry A, 2008. 73(11): p. 975–83.

18. Salumets, A., et al., Epigenetic quantification of immunosenescent CD8(+) TEMRA cells in human blood. Aging Cell, 2022. 21(5): p. e13607.

19. Schmitt, F.A., et al., University of Kentucky Sanders-Brown healthy brain aging volunteers: donor characteristics, procedures and neuropathology. Curr Alzheimer Res, 2012. 9(6): p. 724-33.

20. Abner, E.L., et al., Mild cognitive impairment: statistical models of transition using longitudinal clinical data. Int J Alzheimers Dis, 2012. 2012: p. 291920.

21. Karanth, S., et al., Prevalence and Clinical Phenotype of Quadruple Misfolded Proteins in Older Adults. JAMA Neurol, 2020. 77(10): p. 1299–1307.

22. Nicholas, D.A., et al., Fatty Acid Metabolites Combine with Reduced beta Oxidation to Activate Th17 Inflammation in Human Type 2 Diabetes. Cell Metab, 2019. 30(3): p. 447–461 e5.

23. Winder, Z., et al., Examining the association between blood-based biomarkers and human post mortem neuropathology in the University of Kentucky Alzheimer’s Disease Research Center autopsy cohort. Alzheimers Dement, 2023. 19(1): p. 67-78.

24. Burel, J.G., et al., An Integrated Workflow To Assess Technical and Biological Variability of Cell Population Frequencies in Human Peripheral Blood by Flow Cytometry. J Immunol, 2017. 198(4): p. 1748–1758.

25. Strickland, M., et al., Mitochondrial Dysfunction in CD4+ T Effector Memory RA+ Cells. Biology (Basel), 2023. 12(4).

26. Chen, X., et al., Microglia-mediated T cell infiltration drives neurodegeneration in tauopathy. Nature, 2023. 615(7953): p. 668-677.

27. Laurent, C., et al., Hippocampal T cell infiltration promotes neuroinflammation and cognitive decline in a mouse model of tauopathy. Brain, 2017. 140(1): p. 184–200.

28. Rosenzweig, N., et al., PD-1/PD-L1 checkpoint blockade harnesses monocyte-derived macrophages to combat cognitive impairment in a tauopathy mouse model. Nat Commun, 2019. 10(1): p. 465.

29. Baruch, K., et al., PD-1 immune checkpoint blockade reduces pathology and improves memory in mouse models of Alzheimer’s disease. Nat Med, 2016. 22(2): p. 135–7.

30. Weng, N.P., A.N. Akbar, and J. Goronzy, CD28(-) T cells: their role in the age-associated decline of immune function. Trends Immunol, 2009. 30(7): p. 306–12.

31. Verma, K., et al., Human CD8+ CD57- TEMRA cells: Too young to be called “old”. PLoS One, 2017. 12(5): p. e0177405.

32. Derhovanessian, E., et al., Infection with cytomegalovirus but not herpes simplex virus induces the accumulation of late-differentiated CD4+ and CD8+ T-cells in humans. J Gen Virol, 2011. 92(Pt 12): p. 2746–2756.

33. Wertheimer, A.M., et al., Aging and cytomegalovirus infection differentially and jointly affect distinct circulating T cell subsets in humans. J Immunol, 2014. 192(5): p. 2143–55.

34. Carbone, I., et al., Herpes virus in Alzheimer’s disease: relation to progression of the disease. Neurobiol Aging, 2014. 35(1): p. 122–9.

35. Lovheim, H., et al., Interaction between Cytomegalovirus and Herpes Simplex Virus Type 1 Associated with the Risk of Alzheimer’s Disease Development. J Alzheimers Dis, 2018. 61(3): p. 939–945.

